# Stochastic Wiring of Cell Types Enhances Fitness by Generating Phenotypic Variability

**DOI:** 10.1101/2024.08.07.606541

**Authors:** Divyansha Lachi, Ann Huang, Augustine N. Mavor-Parker, Arna Ghosh, Blake Richards, Anthony Zador

## Abstract

The development of neural connectivity is a crucial biological process that gives rise to diverse brain circuits and behaviors. Neural development is a stochastic process, but this stochasticity is often treated as a nuisance to overcome rather than as a functional advantage. Here we use a computational model, in which connection probabilities between discrete cell types are genetically specified, to investigate the benefits of stochasticity in the development of neural wiring. We show that this model can be viewed as a generalization of a powerful class of artificial neural networks—Bayesian neural networks—where each network parameter is a sample from a distribution. Our results reveal that stochasticity confers a greater benefit in large networks and variable environments, which may explain its role in organisms with larger brains. Surprisingly, we find that the average fitness over a population of agents is higher than a single agent defined by the average connection probability. Our model reveals how developmental stochasticity, by inducing a form of non-heritable phenotypic variability, can increase the probability that at least some individuals will survive in rapidly changing, unpredictable environments. Our results suggest how stochasticity may be an important feature rather than a bug in neural development.

## 1 Introduction

Animals are born with a remarkable array of innate abilities, which enable them to perform essential behaviors from the moment they enter the world, often even in the absence of instructive experience. Colts can stand and walk within hours of birth, spiders can hunt and capture prey, and ducklings can swim almost immediately after hatching—all without parental instruction or extensive trial-and-error. When old field mice, which build long burrows with an emergency exit, are reared from birth by closely related deer mice that build simpler burrows without a second exit, the adopted pups build burrows similar to those of their genetic parents rather than their foster parents (Weber et al., 2013). Even more striking, fish in which all neuronal activity has been blocked during development can swim and respond appropriately to visual stimuli from the moment neuronal activity is restored, demonstrating that these behaviors can develop without any activity-dependent learning (Barabási et al., 2024). These innate abilities, present at birth and developed independently of experience, must be controlled by neural circuitry whose wiring is encoded in the genome and read out by the process of development.

Large brains can contain an enormous number of neurons connected by an even larger number of synapses. Naively, specifying the connectivity matrix of a human brain with ∼10^11^ neurons would require storing connection information for each of the ∼10^14^ synapses, amounting to at least 10^15^ bits of information. In stark contrast, the human genome comprises only about 3 billion base pairs, or roughly 10^9^ bits, revealing a discrepancy of about six orders of magnitude (Wei et al., 2013; Zador, 2019). Thus, even if every nucleotide in the human genome were fully dedicated to optimally and efficiently specifying the precise connections in the brain, the genome would still fall far short of the required information capacity. The limited information capacity of the genome in comparison to the complexity of the connectome, a mismatch that has been referred to as the “genomic bottleneck” (Zador, 2019), implies some form of compression in the developmental specification of brain wiring. The need for such compression underscores the fact that brain development follows orderly rules and is not merely a “look-up table” (Mitchell & Cheney, 2024). This principle is implicit in developmental neuroscience, which seeks to characterize the nature of those rules.

Recent theoretical work has begun to explore how the genome might efficiently encode the connectivity of large brains despite its limited information capacity (Barabási et al., 2023; Koulakov et al., 2021). In the “deterministic genomic bottleneck” (DGB) model (Koulakov et al., 2021), inspired by receptor-ligand based wiring rules, a smaller “genomic” neural network was used to compress the weights of a much larger “phenotypic” artificial neural network model. The model, which was deterministic in the sense that a given model genome reliably and reproducibly generates exactly one connectome, demonstrated excellent “zero-shot” learning—good performance upon initialization, without further training—corresponding to animals’ advanced innate abilities at birth.

Here, we consider the implications of developmental stochasticity (Mitchell, 2018; Ballouz et al., 2023; Linneweber et al., 2020; Vogt, 2015; Koulakov & Tsigankov, 2004; Tsigankov & Koulakov, 2009; Mitchell, 2024)—the fact that a given genome specifies an ensemble of different connectomes—by extending the deterministic genomic bottleneck framework. Building on the “probabilistic skeleton” approach (Stöckl et al., 2021), we propose a model in which the genome encodes the parameters of probability distributions over synaptic strengths between different cell types. Individual connectomes are then sampled from these genetically-encoded distributions. Mathematically, this model can be seen as a generalization of Bayesian neural networks (Blundell et al., 2015), with the genome encoding priors over weights rather than encoding the weights themselves.

We find that this stochastic genomic bottleneck (SGB) model offers several advantages over the DGB model. First, for high levels of compression (i.e., as the number of connections is large relative to the amount of genetic information), the SGB achieves better performance than the DGB, suggesting that stochasticity might be particularly important in larger brains. Second, in simulated physics environments, the SGB model produces individual networks that are specialized to unseen environment niches and body morphologies. Finally, we prove analytically, and confirm empirically, that the average performance of networks sampled from an SGB is higher than the performance of the average network encoded by the genome, providing a population-level evolutionary advantage to stochasticity.

These results suggest that stochasticity may be beneficial for neural development, particularly for animals with large brains that must compress a large amount of connectivity information into a relatively small genome. Stochasticity allows for the generation of diverse networks, some of which may be particularly well-suited to the specific environmental niche of the organism. This work thus provides a theoretical foundation for understanding the role of noise in brain development, and suggests that embracing and harnessing developmental stochasticity may be a key principle in the evolution of complex brains.

## 2 Results

We begin by describing the Stochastic Genomic Bottleneck (SGB) framework in Section 2.1. We then test the SGB framework by applying it to common supervised learning benchmarks in Section 2.2. We focus on what we will refer to as “innate performance” (i.e., performance of the initialized network corresponding to an agent “at birth”, without further training), which in the machine learning literature is often referred to as “zero-shot” learning. Then, we consider the innate performance of the SGB framework in a reinforcement learning setting, on standard continuous control environments (Freeman et al., 2021) in Section 2.3.1, and then in a high-throughput robotics simulator (Makoviychuk et al., 2021) in Section 2.3.2.

### 2.1 Stochastic Genomic Bottleneck Model

In the Deterministic Genomic Bottleneck (DGB), a “genomic” neural network learns to deterministically specify the weight of the synaptic connections between any two neurons in a “phenotypic” network (Koulakov et al., 2021). This phenotypic network is first trained in order to ensure that the genomic network learns connectomes that provide good performance on a task. The genomic network can be thought of as a compressed version of the phenotypic network because it has fewer parameters (refer to Section 4.3.2 for more details).

To study the impact of stochasticity on development, we introduce the Stochastic Genomic Bottleneck (SGB) algorithm. The SGB algorithm, inspired by the probabilistic skeleton of Stöckl et al. (2021), prespecifies neurons in the network as belonging to different cell types and models the connection strength between neurons via cell-type specific connection probabilities. The original model of Stöckl et al. (2021), which employed spiking neurons, was not differentiable and used an evolutionary strategy-based algorithms for optimization. In addition, the probabilistic skeleton had structured connectivity. Here, we reformulate the model to learn the distribution of connection weights between each pair of cell types under the Bayesian framework (Figure 1). Compression occurs as long as the number of cell types is less than the number of neurons. Conceptually, the SGB is a generalization of the standard Bayesian neural network (see Section 4), but rather than specifying a unique weight distribution between every pair of neurons, here instead the weight distribution is shared across all neurons of a given cell type.

**Figure 1:**
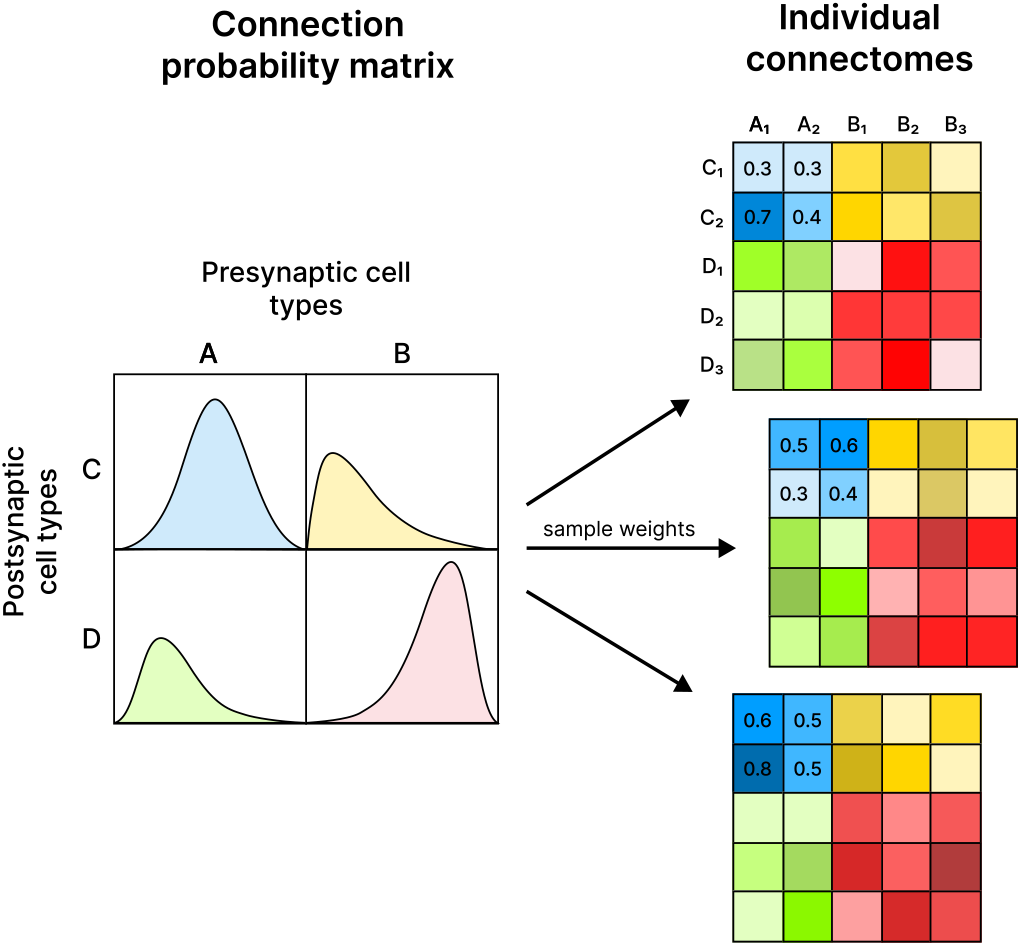
Schematic of the Stochastic Genomic Bottleneck (SGB) The SGB framework uses a low-dimensional parameter set to specify the mean and variance of the synaptic weight distribution between neuronal cell types, allowing individual weights to be sampled stochastically. Unlike the deterministic approach of the DGB, the SGB leverages stochastic sampling to introduce variability in neural wiring. Neurons are grouped into distinct cell types, with their synaptic connections described by a connection probability matrix. This schematic shows the basic setup of generating individual connectomes from the connection probability matrix.

To generate individual connectomes within the SGB framework, we begin with a predefined set of *k* distinct neuronal cell types, denoted as *C* = {*c*_1_, *c*_2_, … *c*_*k*_}. When applying SGB to feedforward layers, each neuronal cell type is either presynaptic or postsynaptic. The synaptic connections between these cell types are therefore described by a two-dimensional connection probability matrix **P**, where each element *P*_*ij*_, indicates the connection probability between the presynaptic neuronal types *i* and postsynaptic neuronal type *j*. Each element *P*_*ij*_ of the connection probability matrix **P** is parameterized by a prior distribution as described in Section 4.3.

Each individual’s connectome **W**_*q*_ is then generated by sampling from the connection probability matrix **P**. The matrix **W**_*q*_ is larger than **P** because it represents the actual synaptic weights between individual neurons, not just between cell types. The generation of **W**_*q*_ involves creating a block structure where each block corresponds to the connection strength between neurons of a specific pair of cell types. This process is formalized as follows:

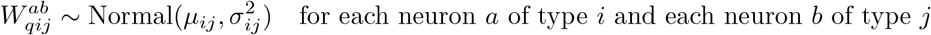

where *a* and *b* index neurons within their respective cell types, and *µ*_*ij*_ and *σ*_*ij*_ are the mean and variance of the Gaussian distributions derived from *P*_*ij*_.

When applied to convolutional layers, we generalize the notion of having distinct cell types to each dimension of the weight tensor. Therefore, the connection probability matrix **P** is four-dimensional, enabling weight sharing along both dimensions of the filters, the input channels, and the output channels. Similarly, we can sample individual connectomes **W**_*q*_ from the connection probability matrix **P**.

The parameters of the SGB are optimized with gradient descent using the reparameterization trick on an evidence lower-bound loss (ELBO) to generate phenotypic networks that perform well at a given task (refer more details in Section 4). In the SGB framework, network complexity and architecture are parametrized by the number of distinct neuronal cell types, *k*. For *k* = 1, the framework reduces to a conventional neural network initialization (except it does not scale weight distributions by the number of input/output connections (Glorot & Bengio, 2010)). Conversely, at the upper limit where *k* equals the number of neurons—effectively treating each neuron as its own distinct cell type—the model transitions to a full standard Bayesian neural network.

### 2.2 Stochastic compression outperforms deterministic compression in supervised learning

We first compared the performance of the SGB to DGB algorithms on two well-studied supervised learning tasks, the MNIST and CIFAR image classification benchmarks (Section 2.1). As noted above, both the DGB and SGB seek to compress a phenotypic network into a smaller set of parameters, but whereas the DGB deterministically sets the synaptic weights in the phenotypic network, in the SGB the parameters specify probability distributions over the synaptic weights of the phenotypic network. We evaluate the performance of these models under varying levels of compression, defined as the ratio between the number of parameters in a standard uncompressed network and an equivalent compressed network Figure 2.2. For both models, “innate performance” (defined as performance of the SGB- or DGB-initialized model on held-out test data before any further training data are presented) is plotted as a function of compression. As expected, performance drops monotonically with increasing compression for both models, analogous to how lossy image compression techniques such as jpeg result in blurrier reconstructions as compression increases.

**Figure 2:**
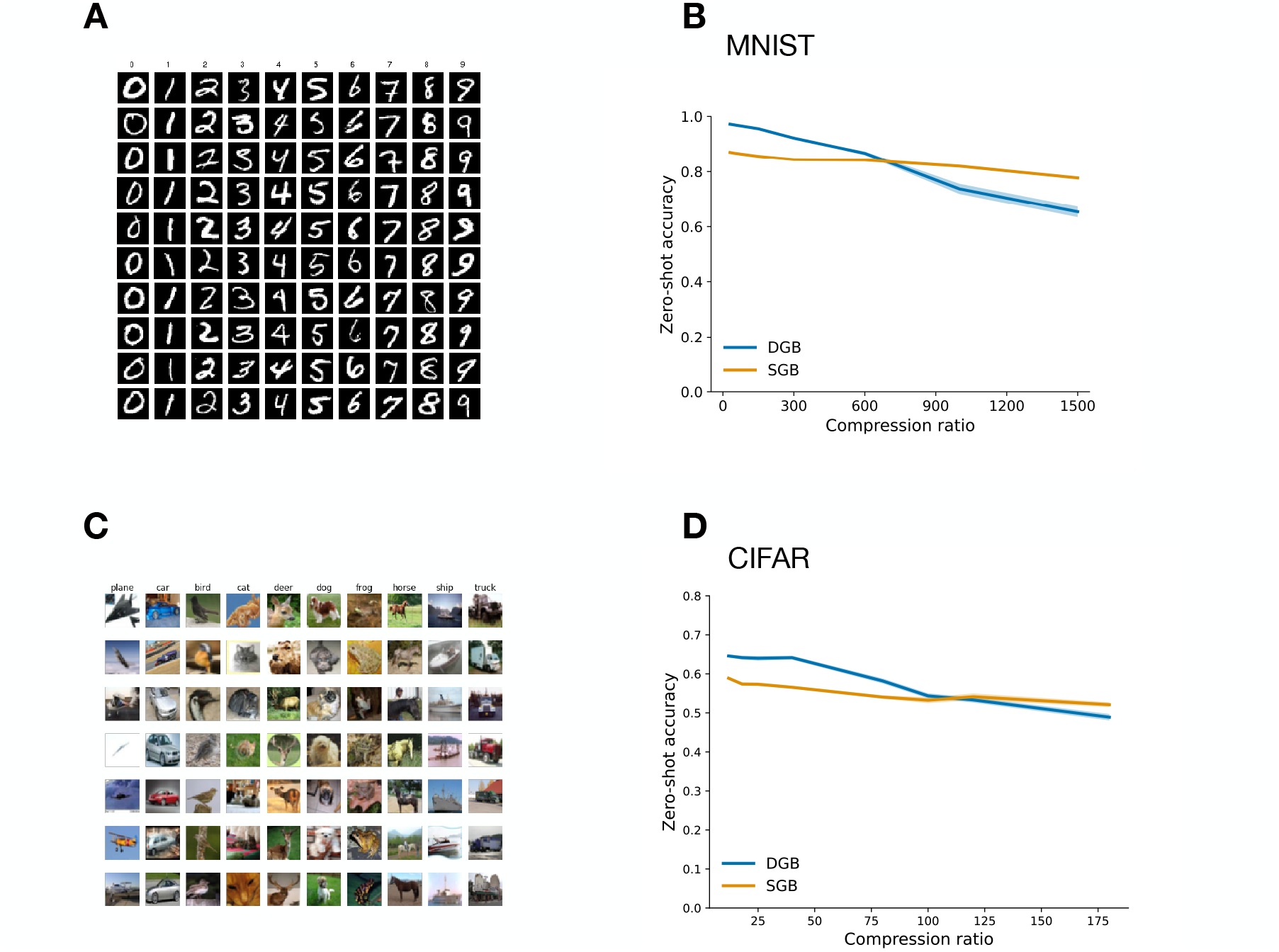
Stochastic genomic gottleneck (SGB) leads to better innate performance at high compression compared to the deterministic genomic bottleneck (DGB). **(A)** Examples of MNIST dataset of handwritten digits. **(B)** Innate performance on MNIST, measured by the zero-shot classification accuracy, for both the DGB and SGB models across various levels of compression. Solid lines represent the mean zero-shot accuracy across five seeds with shaded areas representing the standard error of the mean. **(C)** Examples of CIFAR dataset of images. **(D)** Innate performance on CIFAR for both DGB and SGB model for various levels of compression across five seeds.

Comparing the two models, we find that for small compression levels the DGB performs better, but at higher levels of compression the SGB outperforms the DGB on both MNIST and CIFAR. Because the DGB is a special case of the SGB, in which the standard deviation of the weight distribution is zero, it might seem surprising that the DGB could ever outperform the SGB. However, the KL divergence penalty imposed on the posterior in the SGB prevents the standard deviation from reaching zero, ensuring the posterior remains a valid probabilistic distribution (refer to equation 1). If it did reach zero, the KL divergence would become infinite. This accounts for the observed performance differences at lower compression levels.

Furthermore, the SGB maintains remarkably stable performance across a wide range of compression levels, demonstrating its robustness and efficiency in handling high compression. These results suggest that stochastic developmental processes may be particularly beneficial for organisms with large brains, which face a greater challenge in genetically encoding their vast neural connectivity patterns (Zador, 2019).

### 2.3 SGB for Reinforcement Learning

We next tested the SGB framework on reinforcement learning tasks, which are more relevant to the kinds of learning and decision-making problems faced by animals in their natural environments. In contrast to supervised learning, where the goal is to learn a mapping from inputs to outputs based on a labeled dataset, reinforcement learning involves learning to make a sequence of actions in an environment to maximize a cumulative reward signal (Sutton & Barto, 2018). This trial-and-error process, guided by rewards, more closely resembles how animals learn from experience to navigate their world and acquire complex behaviors.

We used reinforcement learning tasks inspired by the motor control challenges faced by animals. Just as organisms must learn to efficiently control their bodies to navigate environments and acquire resources, our RL agents must learn to map sensory inputs to motor outputs to maximize reward. This allows us to study how an agent’s learning and performance are shaped by its embodiment and environment (Pfeifer & Bongard, 2006), key factors in both biological and artificial intelligence.

We test SGB on continuous control tasks in a simulated physics environment, BRAX (Freeman et al., 2021), which loosely mimics the motor control challenges of legged animals. In Section 2.3.1, we benchmark SGB on the Ant and Halfcheetah environments from Freeman et al. (2021). In Section 2.3.2, we experiment with controlling the Ant agent with different body morphologies, and controlling a simulated quadruped ANYmal robot (Hutter et al., 2016) on different terrains.

For our reinforcement learning experiments, we employ Proximal Policy Optimization (PPO) (Schulman et al., 2017), a standard on-policy actor-critic algorithm that offers a good trade-off between sample efficiency and wall-clock training time. We use SGB layers throughout the agent’s policy and value networks, except for the output layers which remain deterministic. Full architectural details and hyperparameters are provided in Appendix 4.5. We train SGB agents with varying numbers of cell types, as shown in Figure 3, and measure their performance as a fraction of the return achieved by an uncompressed baseline agent. For the Brax experiments, the baseline agent has a hidden layer width of 128, so an SGB agent with 128 cell types experiences no compression. For the ANYmal experiments, the SGB agent has 256 cell types compared to the baseline’s hidden layer width of 1024.

**Figure 3:**
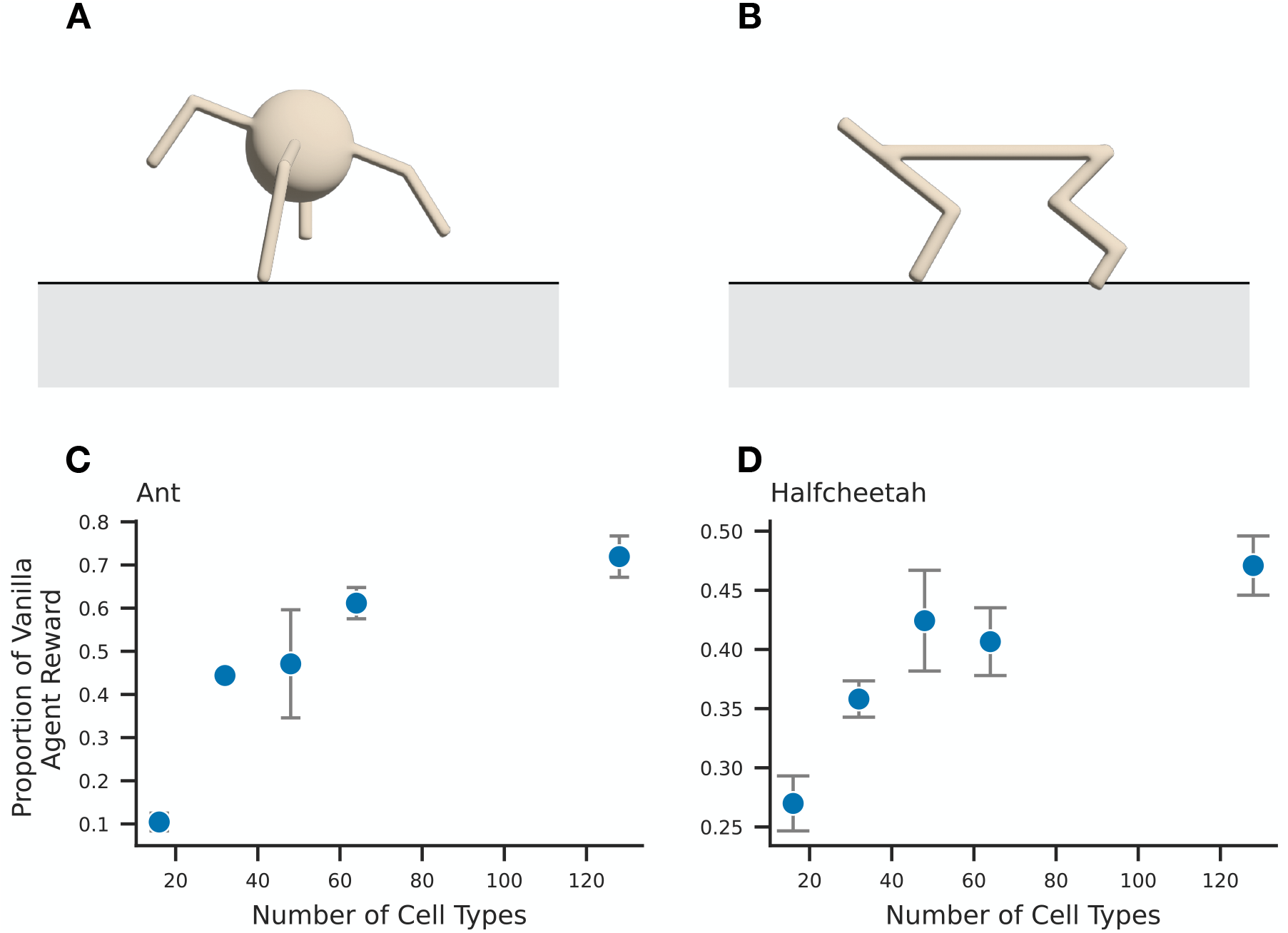
SGB endows reinforcement learning agents with innate motor control capacities. **(A)** Illustration of the Ant reinforcement learning environment where the network learns to coordinate the four legs of the ant to move forward as fast as possible. **(B)** Illustration of Halfcheetah environment where the goal is to apply appropriate torques on the joints to make the halfcheetah run as fast as possible. **(C)** Zero-shot performance as a function of different prespecified number of cell types in the network. Here, zero-shot performance is measured by the reward obtained by the compressed stochastic networks (the SGB models) normalized by that of networks without compression. Error bars represent standard error of the mean across five seeds. **(D)** Similar to (C) but for the HalfCheetah environment.

#### 2.3.1 Stochastic compression on motor control tasks

In both the Ant and Halfcheetah tasks, the goal for the corresponding agent is to learn to move its body forward, while minimizing constraints such as the energy expended. The state space of both tasks consists of the positions and velocities of the agent’s limbs, and the action space is a continuous vector of forces it can apply to its different limbs. The performance of the Ant and Halfcheetah agents as a proportion of a vanilla PPO agent (non-compressed, deterministic) is shown in Figure 3. When the hidden layer width of the SGB agent is equal to that of the vanilla agent, the SGB agent is able to achieve a significant proportion of the return achieved by the vanilla agent (≈75% for Ant and ≈45% for Halfcheetah). As the number of cell types decreases, the performance of the SGB decreases gradually until the region of 16 cell types, where the SGB performance falls off quickly, indicating that it is in this region where the SGB network does not have enough expressive capacity to represent a reasonable policy for the SGB. These results suggest that there is a minimal number of cell types needed to endow a network with the needed flexibility.

#### 2.3.2 SGB induces population-level diversity in behavioral capacity

##### Ant motor control with varied leg lengths

Next, we sought to examine the influence of developmental stochasticity on the phenotypic diversity and adaptability of reinforcement learning agents to variations in body morphologies. We optimized the SGB on the Ant environment and sampled 100 agents generated by the SGB. These networks were tested for their innate (zero-shot) performance within the same environment and on a modified version of the Ant environment where the four legs of the agent were lengthened Figure 4B. Although on long time scales it seems plausible that brains and bodies co-evolve, on short time scales the brain specified by a given genetic program must learn to control whatever body happens to develop, which can be shaped by forces in an animal’s lifetime (e.g. the availability of nutrients for growth).

**Figure 4:**
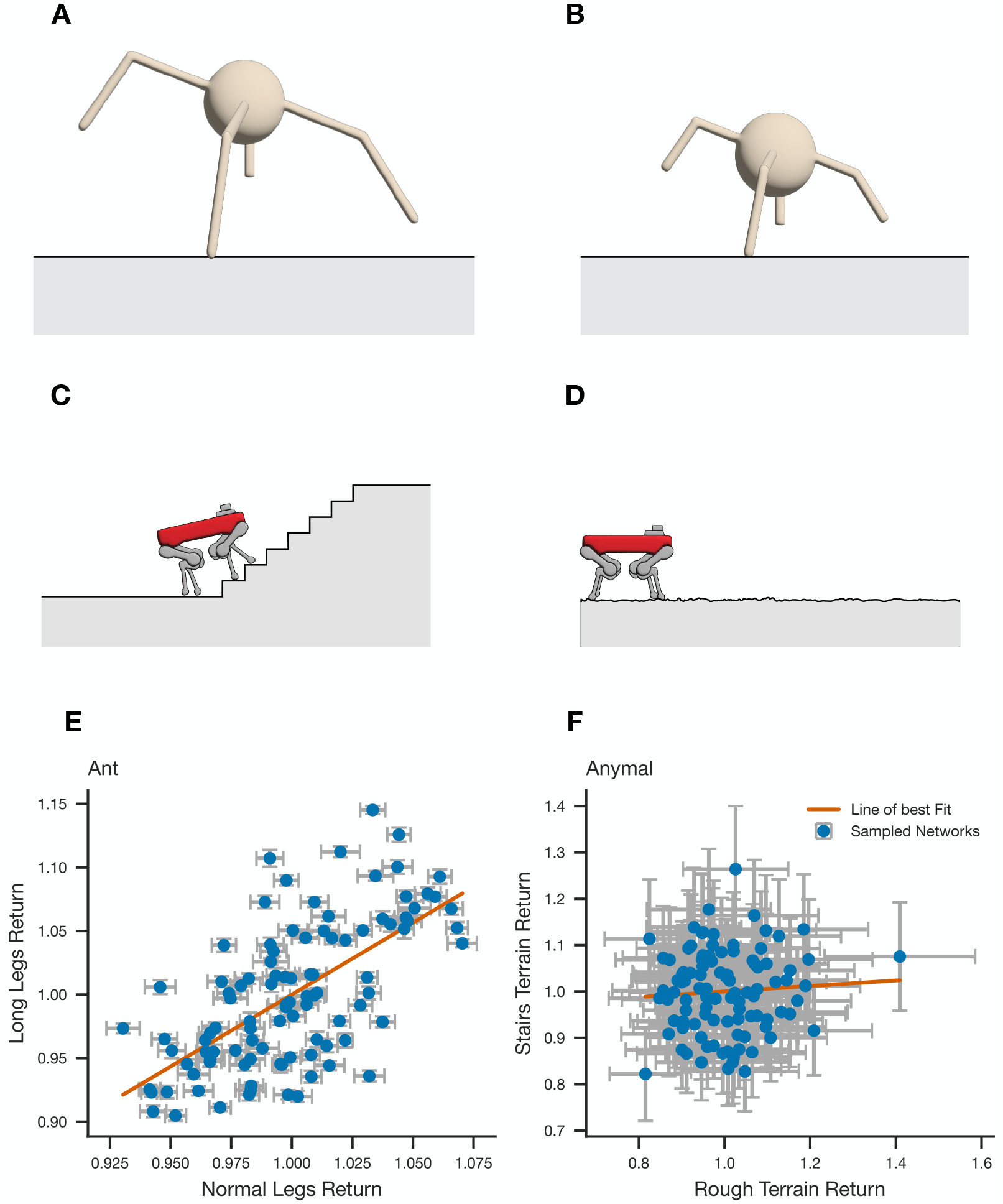
SGB induces phenotypic diversity for different body morphologies and environmental terrains. **(A)** Illustration of the Ant environment with long legs, contrasted with **(B)** the Ant environment with normal legs. **(C)** Illustration showing Anymal robot^1^ on the stairs terrain versus **(D)** the rough terrain. **(E)** Innate performance of sampled networks for the long leg versus normal leg Ant environment. The SGB is first evolved on the normal leg environment. After the SGB has been optimized, we sampled networks from the trained SGB and tested their zero-shot performance without any learning on both environments. Each dot is one sampled network from the evolved SGB tested on the two environments. The networks’ returns are positively correlated on the two body morphologies (Pearson’s *r* = 0.64, *p <* 0.001), while also exhibiting phenotypic variability in their capacities for each body morphology. The error bars on each dot represent the standard error of the mean in the return (the reward) obtained by the sampled network across multiple episodes on the corresponding environment. **(F)** Similar to (E), where the SGB was evolved on only the stair terrain, while the subsequently sampled networks were tested for their zero-shot performance on both the stairs and the rough terrains. The sampled networks exhibit substantial phenotypic diversity such that on each environmental terrain, there are individual networks performing notable better than the rest of the population. There is no significant correlation between a network’s performance on rough terrain and its performance on stairs (Pearson’s *r* = 0.065, *p* = 0.52).

Figure 4E shows a scatterplot of the performance of each network controlling a standard body (normal legs) vs a body with long legs. Strikingly, there is considerable scatter around the mean for both conditions, indicating that different networks drawn from the same distribution can achieve very different levels of performance with different body morphologies. Thus, developmental stochasticity can lead to marked phenotypic diversity of behavioral capacities over the population. Furthermore, performance on the two conditions are positively correlated (Pearson’s r = 0.64, p < 0.001), suggesting that certain “athletic” neural phenotypes are intrinsically better at performing the locomotion task, regardless of variations in body morphology.

##### ANYmal Robot Control on Varied Terrains

In nature, animals must navigate a wide variety of terrains, from flat plains to rocky hillsides. From an evolutionary perspective, this diversity of environments may favor the development of specialized individuals within a population, each adapted to a particular niche. We hypothesize that stochastic development, by introducing variability in neural wiring, could be a mechanism for generating this specialization. To test this idea in a controlled setting, we use the ANYmal robot environment from Rudin et al. (2021), which challenges a reinforcement learning agent to control a quadruped robot in traversing different terrains, including rough terrain and stairs. The agent must learn to map sensory inputs (the state space, containing information about the terrain and past actions) to motor outputs (the action space, dictating the torques applied to the robot’s joints) in order to navigate each terrain successfully.

We train the SGB on all terrains simultaneously, mimicking the diversity of environments encountered in nature. Then, at test time, we evaluate the innate performance of 100 networks sampled from the SGB on the “rough” and “stairs” terrains separately. Strikingly, we find that the sampled networks show considerable individual differences in their ability to navigate the two terrains. Some networks excel at traversing rough terrain but struggle with stairs, while others show the opposite specialization. Quantitatively, we find no significant correlation between a network’s performance on rough terrain and its performance on stairs (Pearson’s *r* = 0.065, *p* = 0.52), unlike the strong positive correlation between long-leg and short-leg performance in the Ant task (Section 2.3.2). This result suggests that developmental noise can indeed lead to specialization to environmental niches, with different individuals within a population being better adapted to different niches.

### 2.4 Stochastic sampling gives better average performance

To investigate the possibility that the stochasticity of brain development may actually be advantageous, we compared the performance of networks created via stochastic sampling to that of a “mean network”, i.e. where the synaptic weights are simply set to be the means of the specified distributions, which represents the limit case of sampling with zero variance. We conducted this comparison in the reinforcement learning setting, using the Ant environment, and evaluated the mean performance over 100 rollouts for both the sampled networks and the mean network.

Strikingly, we found that the mean performance of a population of stochastically sampled networks exceeds the performance of the mean network. We found that the mean performance over 100 rollouts was substantially higher for the sampled networks than for the mean network (*t*(99) = 12, *p <* 0.001) Figure 5). Assuming a monotonic relationship between the task-specific loss function and performance (where lower loss implies higher performance), these results can be summarized as follows: the loss of the mean network is greater than the mean loss of the sampled networks. According to Jensen’s inequality (Jensen, 1906), this outcome suggests a concave loss function. In Appendix (A.2), we provide a mathematical proof demonstrating how gradient-based optimization leads to sampling networks over a concave slice of the loss landscape. The proof offers a mechanistic explanation for the seemingly counterintuitive result: A diverse population of brains, each wired slightly differently due to developmental noise, collectively outperforms a population of identical brains wired according to the mean connectivity. Thus, at the population level, stochasticity yields better performance.

**Figure 5:**
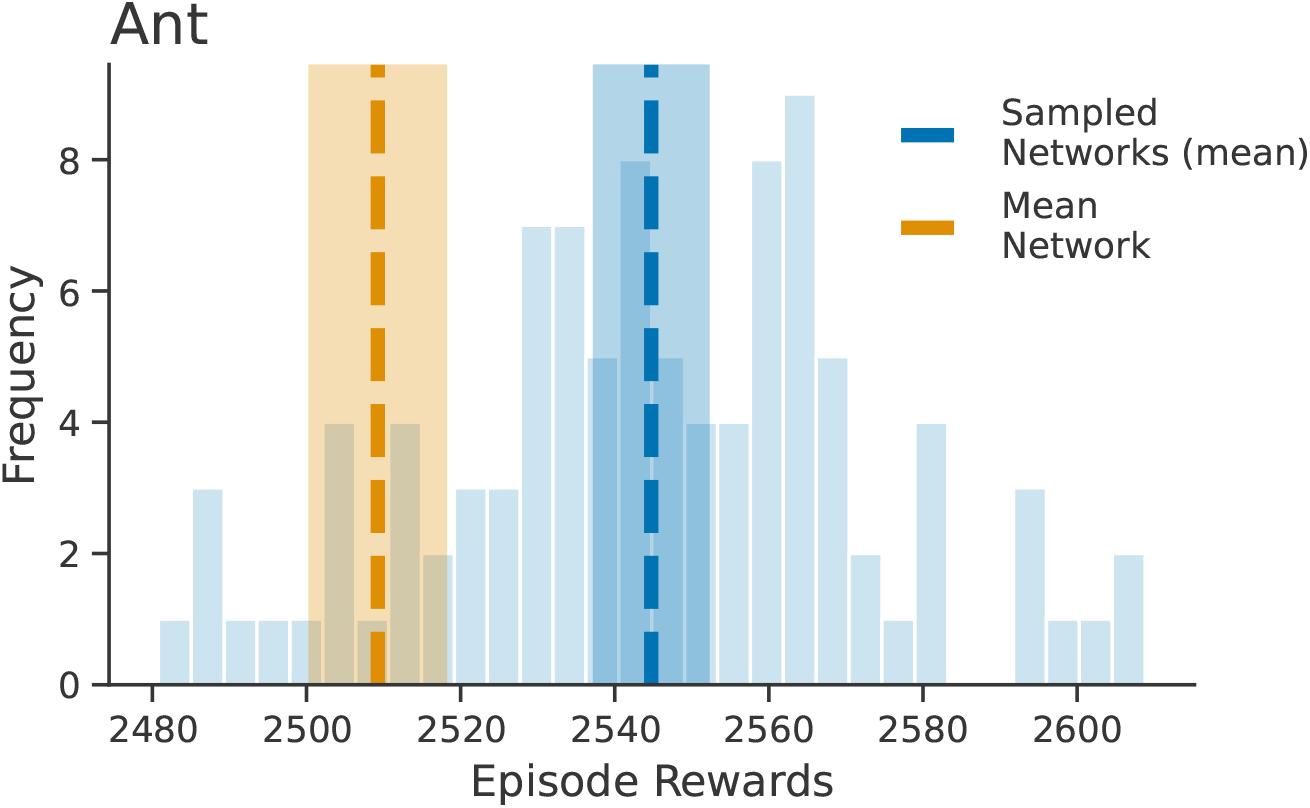
Stochastic sampling gives better average performance across a population of sampled networks. Zero-shot performance of sampled networks and mean network with 64 cell types on the Ant environment. Shaded region indicates 99% confidence intervals.

**Figure 6:**
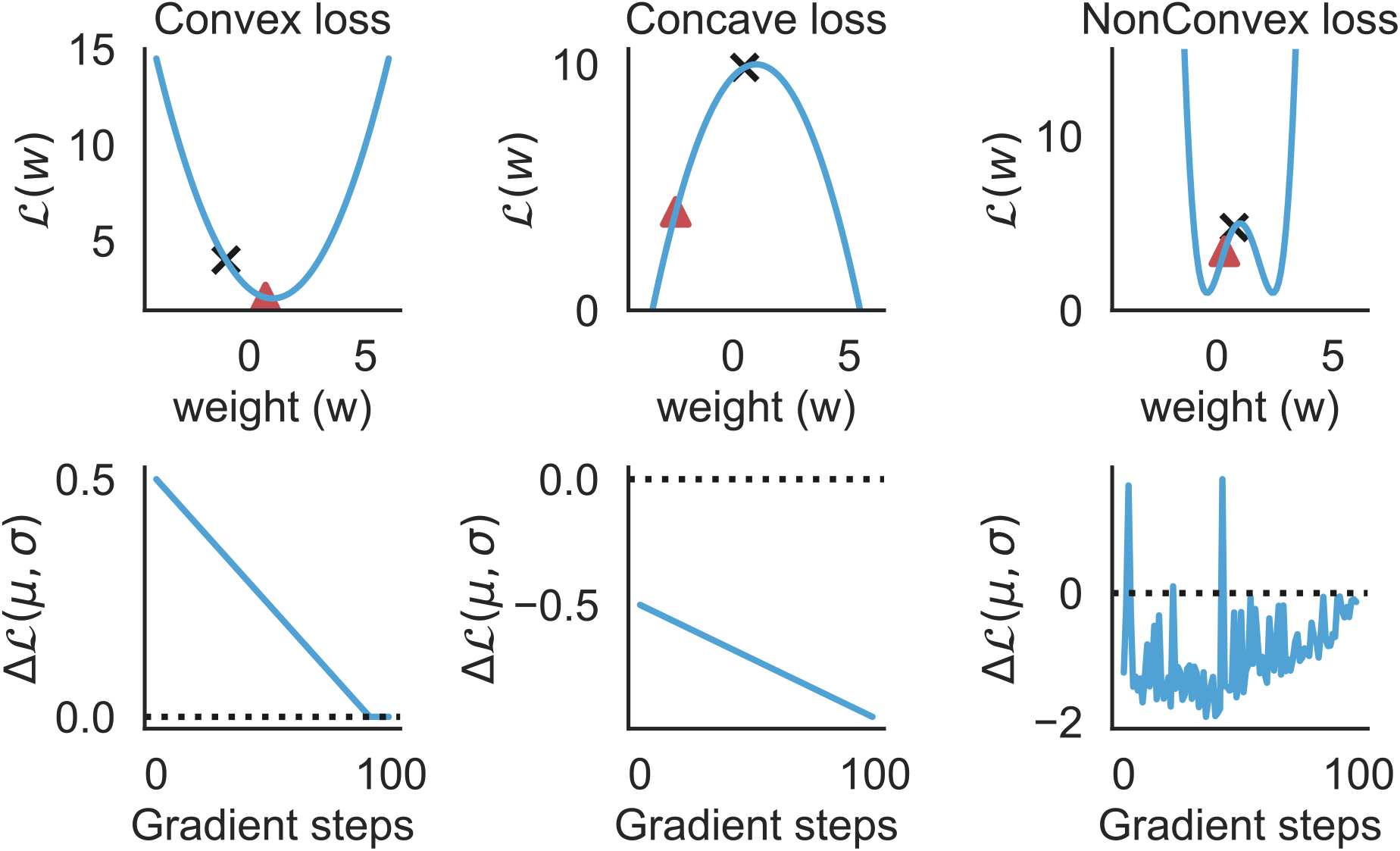
Empirical verification of subsection A.5 in toy 1D settings with Convex (left), Concave (middle) and NonConvex (right) loss functions. The black cross and the red triangle indicate the initial and final (after 100 gradient steps) parameter values of the mean network, *µ*. Note that Δℒ (*µ, σ*) is always ≥ 0 for the Convex loss landscape and ≤ 0 for the Concave loss landscape. For the NonConvex loss landscape, Δℒ (*µ, σ*) depends on the location of *µ* in the parameter space.

**Figure 7:**
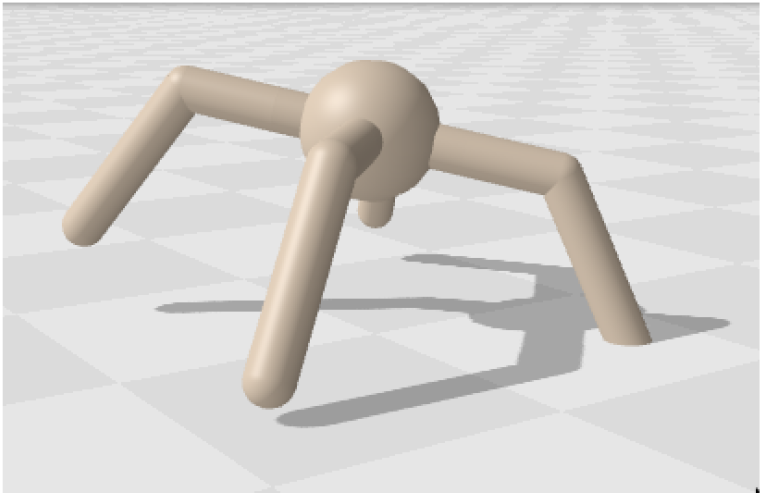
Ant environment in the Brax simulator.

**Figure 8:**
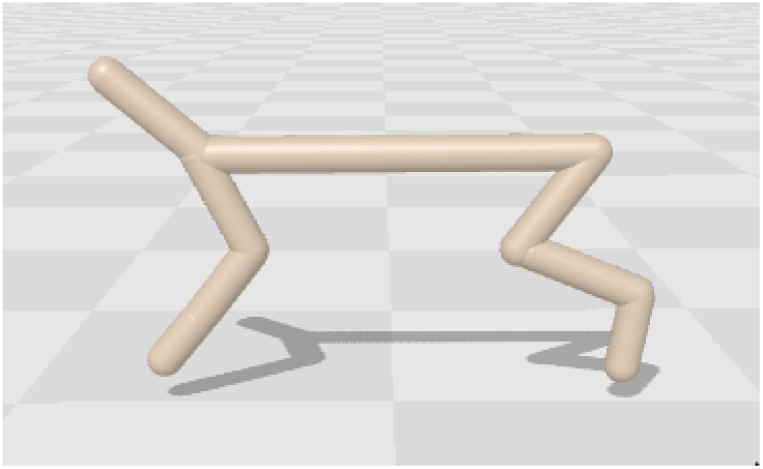
Cheetah environment in the Brax simulator.

**Figure 9:**
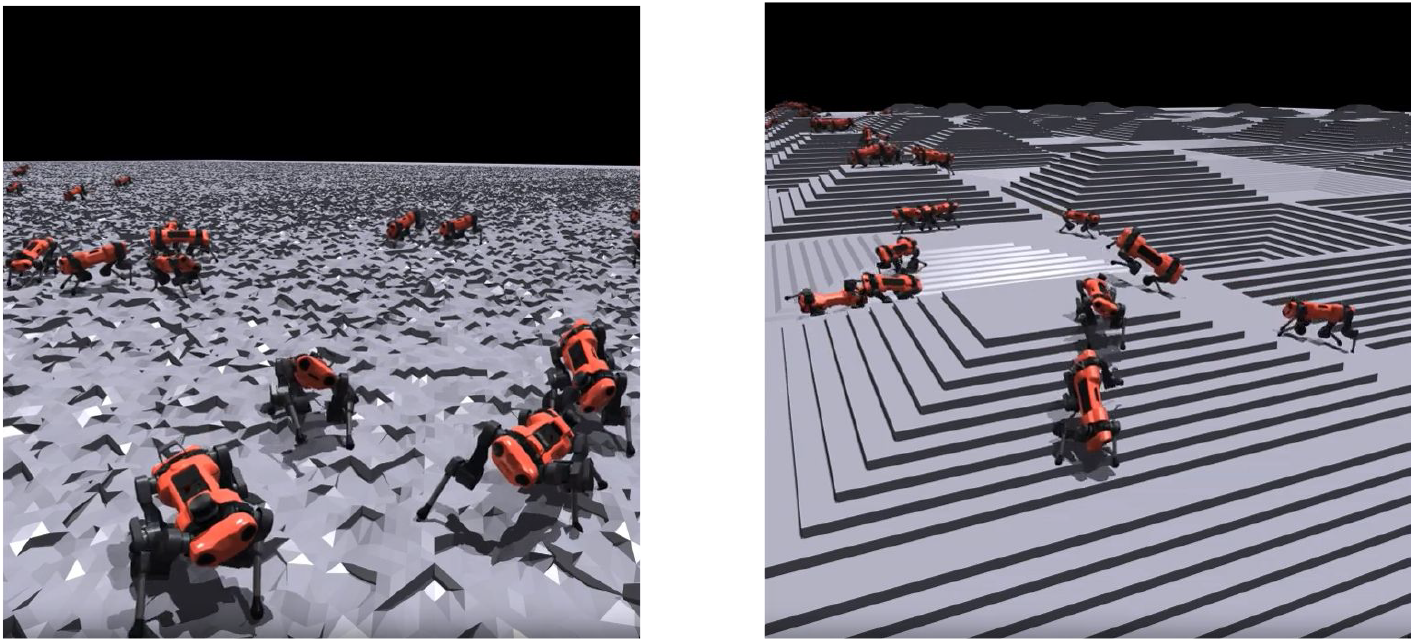
Multiple instances of the Anymal robot in the Brax simulator. The left image shows robots on rough terrain, while the right image shows them on a stair terrain. Different agents represent separate simulations with no interaction between agents.

These results suggest that the stochasticity of brain development may not be merely a bug, but rather a feature that evolution has harnessed to enhance behavioral adaptability. A population of diverse brains, collectively exploring a wider range of cognitive and behavioral strategies, may be more resilient to environmental challenges than a phenotypically identical population. This diversification may be especially advantageous in the face of compression.

## 3 Discussion

In this study, we introduced the Stochastic Genomic Bottleneck (SGB) model, a differentiable algorithm that incorporates stochasticity into the process of generating neural network architectures from compressed genomic representations (Stöckl et al., 2021). Inspired by the noisy nature of biological development (Mitchell, 2018; Ballouz et al., 2023; Linneweber et al., 2020), our model suggests that stochastic rules provide a powerful approach to overcoming the limited information contained in the genome, resulting in the generation of diverse and complex neural connectomes. Our model can be seen as a generalization of Bayesian neural networks and offers a new perspective on how brains can efficiently develop from genetic information.

Through experiments on both supervised learning and reinforcement learning tasks, we demonstrated that SGB maintains stability across a wide range of compression levels and significantly outperforms deterministic models in high compression settings. We find that SGB excels in large networks and variable environments, exhibiting superior adaptability and robustness due to the incorporation of stochasticity. Furthermore, we provide analytical proofs and empirical validation showing that the mean performance of connectomes sampled through our stochastic model surpasses the performance of the average network generated without stochasticity in complex loss landscapes. This indicates that stochastic sampling around the mean provides an advantage over generating each synaptic weight precisely according to a deterministic rule.

Our model draws an analogy between the processes of evolution and development. The optimization of cell type connectivities in our model represents the role of evolution in shaping the genetic blueprint for brain development. For computational efficiency, we have constructed our model to be differentiable, which greatly facilitates navigation in high-dimensional spaces. However, it is important to note that at the level of an individual, Darwinian evolution is not differentiable: the specific genetic mutations and recombinations that appear in the offspring do not benefit from the life experiences of the parents. Thus, we are not positing that our model is an accurate reflection of evolution and neural development. Rather, it provides a proof-of-concept that stochastic development can be functionally useful.

The phenotypic variability discussed in our study differs from traditional sources of phenotypic variability, which arise from genotypic variability and environmental factors (nature and nurture). The importance of stochasticity (chance) as a third source of phenotypic variability is increasingly recognized (Mitchell, 2018). While phenotypic variability arising from genotypic variability is heritable, variability resulting from stochasticity is not. Growing evidence supports the significance of stochastically induced phenotypic variability. For example, in the nine-banded armadillo, a species which uniquely among mammals reproduces with genetically identical quadruplets, minor stochastic variations in gene expression during early development have lasting impacts. This highlights the profound influence of developmental noise on individual diversity beyond genetic and environmental determinants (Ballouz et al., 2023). In Drosophila melanogaster, nonheritable, stochastic variations in brain wiring within the dorsal cluster neurons (DCN) contribute to individual differences in object orientation behavior, demonstrating a direct link between brain development noise and behavioral variation. Furthermore, in neurodevelopmental disorders like schizophrenia and autism concordance can be as high as 50% (Imamura et al., 2020), raising the possibility that early stochastic development may play a role in the remaining variation.

In engineering, noise is often considered a nuisance, and great efforts are made to manufacture parts with high precision. However, our findings suggest that in some biological contexts, noise may actually be advantageous. The incorporation of stochasticity in neural development can lead to increased adaptability and robustness, especially in large networks and variable environments.

Our work builds on a long tradition of computational models exploring how patterned neural connectivity emerges through development. Early pioneers like Turing proposed models of how simple local interactions could give rise to structured patterns in biological systems (Turing, 1952). In the domain of neural development, Hebbian learning rules can explain the emergence of center-surround receptive fields and ocular dominance columns in the visual system (Linsker, 1988; Miller et al., 1989; Miller & MacKay, 1994). More recently, researchers have explored how structured connectivity can arise through various mechanisms, including synaptic plasticity (Clopath et al., 2010), axon guidance (Goodhill, 2007), and spontaneous activity (Albert et al., 2008). However, these models typically focus on recapitulating observed connectivity patterns rather than directly optimizing for computational function. In contrast, our approach starts from the assumption that the network must perform a specific task and asks how this computational goal constrains the developmental process. By directly optimizing for performance, our model provides a complementary perspective on how evolution and development might shape neural connectivity to support adaptive behavior.

The significance of our findings goes beyond biological inspiration. They suggest new directions for developing AI models that are more flexible and capable of dealing with uncertain environments. For example, ANN researchers could first optimize an SGB in a generic environment, sample individual networks with good zero-shot performance from the SGB, and then train those individual networks on various tasks with much less data than would be required to train them from scratch. This could prove particularly advantageous in reinforcement learning settings where data collection can be expensive and time-consuming.

Our results highlight the power of incorporating randomness into the development of neural networks, offering a more nuanced approach to bridging genetics and brain structure. Our work not only advances our understanding of biological processes but also opens up new possibilities for creating more adaptable and resilient artificial intelligence systems. By embracing the inherent stochasticity in developmental processes, we hope to gain insights not only into the evolution of natural intelligence but also the design of artificial ones.

## 4 Methods

### 4.1 Preliminaries

First, we describe the model in the supervised learning setting. Using the notation of (Goodfellow et al., 2016), let **X** = {**x**_1_, **x**_2_, …, **x**_*M*_} denote our data and **Y** = {**y**_1_, **y**_2_, …, **y**_*M*_} be the targets. Typically, deep neural networks train on a maximum likelihood loss, which aims to maximise the likelihood of the targets given the input data and neural network parameters

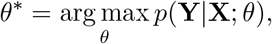

where *θ*^∗^ is the set of optimal parameters (Goodfellow et al., 2016; Blundell et al., 2015). In the case of the stochastic genomic bottleneck, we instead aim to learn a posterior distribution over weights with approximate Bayesian inference.

### 4.2 Bayesian Neural Networks

Bayesian neural networks (BNNs) have gained significant attention in the machine learning community due to their ability to capture uncertainty in model predictions by treating the weights of a neural network as probability distributions rather than fixed values (Blundell et al., 2015; Wilson & Izmailov, 2022; Ghahramani, 2015; Kendall & Gal, 2017). This allows BNNs to express the confidence of their predictions, which is particularly valuable in decision-making scenarios where the cost of an incorrect prediction is high (Kendall & Gal, 2017; Leibig et al., 2017; Kahn et al., 2017). Furthermore, uncertainty quantification makes BNNs more robust to overfitting, enables them to generalize better to new data, and provides a principled way to incorporate prior knowledge into the model, allowing for more efficient learning and better performance in data-scarce settings Blun-dell et al. (2015); Fortunato et al. (2017); Louizos & Welling (2017). The Bayesian framework also facilitates model comparison and selection, as it naturally penalizes overly complex models through the marginal likelihood (MacKay, 1992). These advantages have led to the application of BNNs in various domains, such as healthcare, finance, and autonomous systems, where reliable uncertainty estimates and robust decision-making are crucial (Kahn et al., 2017; Leibig et al., 2017).

Instead of learning a single scalar value for each weight and bias parameter, BNNs aim to learn a posterior distribution over weights by employing Bayes rule, meaning weight matrices are sampled from an underlying distribution parametrized by *θ*: **W**∼ *P* (**W** |*θ*) (Blundell et al., 2015). However, when the input data **X** is high-dimensional, computing the full posterior distribution over weights becomes computationally expensive. As a result, practitioners typically learn an approximate posterior with “Bayes by Backprop”, where variational inference is performed by maximizing the ELBO (Blundell et al., 2015):

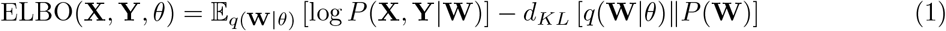

where **W** refers to the set of weights of the BNN, and *q*(**W** | *θ*) is the distribution that approximates the true posterior. Intuitively, maximizing the ELBO requires optimizing the posterior to match the prior distribution *P* (**W**) while also satisfying the likelihood of data under network weights. In practice, each conventional neural network weight is represented by a mean and a variance parameter. The ELBO is then maximized by averaging losses over multiple samples from the weight distributions using the reparameterization trick (Kingma & Welling, 2013), which allows for backpropagating gradients through to the underlying distribution parameters when stochastically sampling network weights.

The SGB framework combines the idea of Bayesian neural networks with the genomic bottleneck (Koulakov et al., 2021). The genomic bottleneck builds upon hyper-networks (Ha et al., 2017), which are (relatively) small networks that are used to generate the weights of a larger network. Krueger et al. (2017) also combine the ideas of Bayesian neural networks and hyper-networks, but generates a particular set of parameters instead of generating all parameters of the network (as we do in SGB).

### 4.3 Parameterization of Weight Distributions

Instead of defining the distribution of connection strength between each pair of neurons individually, SGB groups neurons into different cell types, and defines a distribution over connections between different cell types. As is common in Bayesian deep learning, we also assume zero covariance between the cell type connection distributions, meaning that the variations in connection strength between different cell types are assumed to be independent of each other. So, instead of defining a prior of the whole weight matrix, the stochastic genomic bottleneck defines distributions over connections between different cell types, whose joint probability can be expressed as the following.

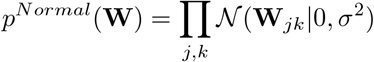

Where **W**_*jk*_ represents the weight between *j*^*th*^ and *k*^*th*^ cell type within the network. Furthermore, instead of using a normal distribution, we follow Blundell et al. (2015) and use a Scale Mixture Prior (SMP). The SMP is characterized by two Gaussian components with zero mean but two distinct variances 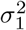 and 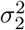, and a mixing coefficient *π*. The SMP is formally defined as follows:

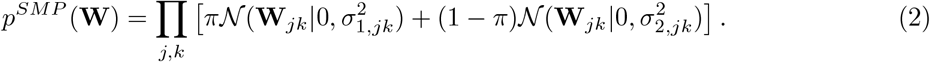

The variances *σ*_1_ and *σ*_2_ allow the model to capture a broad range of weight magnitudes. In our experiments, the initialization of the mean parameters is set to zero, reflecting the assumption of no a priori preferred direction in the weight space. For the variance parameters, 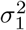 is initialized to capture the larger scale behavior of weights, whereas 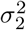 is set several orders of magnitude smaller to encourage sparsity (Blundell et al., 2015). The mixing coefficient *π* is typically initialized to favor the sparser Gaussian component (i.e closer to zero), thus initializing the network in a state that promotes regularization.

To obtain the weight matrix used for a forward pass of a given SGB layer, multiple samples are taken from each distribution *P* (**W**_*jk*_), where *jk* indexes the connection between two cell types. Then, these sampled connections are assigned to entries within a weight matrix that represent a scalar connection between two cell types in two successive layers (as shown in figure 1). Connections between two cell types are represented multiple times within a given weight matrix as there are multiple instantiations of the same cell type in each SGB layer—hence the compressive nature of SGB and the need for multiple samples of *p*(**W**_*jk*_). Gradient are backpropagated, again with reparametrization trick, through the stochastic sampling operations used to obtain weights between two given cell types.

#### 4.3.1 Supervised Learning Loss Function

To optimize the SGB to perform image classification, we employ a standard cross entropy loss for the log-likelihood term in equation 1, (see e.g. Murphy (2012, p. 57)), and then compute the KL divergence between the SMP prior and its learned values for the first term in equation 1, resulting in the following loss function.

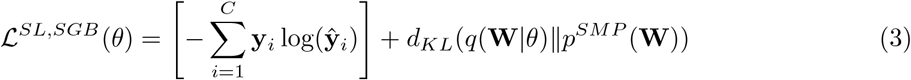

where C is the number of classes in the classification problem, **y** is a one-hot encoded vector describing the ground truth class and **ŷ** is the predicted probabilities for each class. For each gradient update, we aggregate gradients over multiple samples of the weight distribution whose loses are computed across a minibatch of multiple input-output pairs.

#### 4.3.2 Deterministic Genomic Bottleneck

The genomic bottleneck framework, as proposed by Koulakov and colleagues (Koulakov et al., 2021), introduced a method to train ANN’s that have good innate performance. It is inspired by the biological constraint that the genome cannot explicitly encode the vast number of synaptic connections in the brain. Instead, the genome must rely on a compact set of instructions or developmental rules to construct the neural connectome (Zador, 2019). They formulated this problem in terms of lossy compression of a weight matrix of a traditional ANN. The genomic bottleneck approach introduced two distinct networks: the phenotype network (p-net) and a genomic network (g-net). The p-net represents the functional ANN as it would be after traditional training, embodying the learned tasks through its weights and connections. The g-net is tasked with compressing this p-net, which aims to encode p-network’s connectivity and functionality. The authors used an iterative algorithm to learn a g-net that can generate a p-net that has good innate performance. First they train the p-net using standard gradient descent based approach up to some level of performance. Subsequently, the trained weight matrix of the p-net is used to train the weights of the g-net. This process is repeated until the g-net converges to a stable solution i.e it can effectively generate p-nets that have good innate performance and are compressible. This is the training process we use in the experimental sections when reporting DGB performance.

#### 4.3.3 Reinforcement Learning Loss Function

For our reinforcement learning experiments, we use the PPO loss function within the SGB framework. PPO is an actor-critic architecture consisting of two networks (one actor that selects actions and one critic that judges the value of states). The PPO loss function aims to select actions that maximise value. However, PPO is constrained in the size of the updates it can make in order to stabilise training. There is also an entropy term, which encourages exploration during policy learning. PPO works by collecting trajectories from parallel actors and then updating its networks weights with epochs of training on the collected data. The standard PPO objective from Schulman et al. (2017) is as follows:

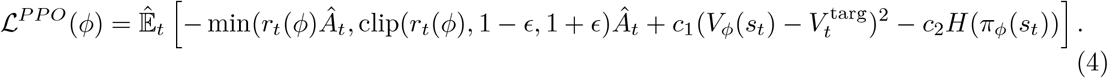

Where we use *ϕ* to represent the trainable weight parameters of a standard deterministic actor-critic network (to avoid confusion with the trainable parameters of a SGB network which we label as *θ*), *r*_*t*_(*ϕ*) is the ratio of the new policy to the old policy, *ϵ* is a clipping parameter that limits the size of gradient updates to improve stability, *Â*_*t*_ is the estimated advantage—which computes the difference between the experienced (bootstrapped) value of the current trajectory compared to its predicted value, *V*_*ϕ*_ is a value function that predicts the value of a given state, 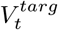 is the bootstrapped target for the value function, *H* computes the entropy of the network policy *π*_*ϕ*_ and *c*_1_ and *c*_2_ are constants that control relative contribution of the different terms to the loss function. Note, we use PPO in environments with continuous action spaces, meaning we use the standard approach of predict parameters of an action distribution, which is then sampled to get the action used by the agent.

In order to adapt PPO to the SGB framework, we use a SGB actor and critic parametrised by *θ* and we let the traditional PPO loss function represent the log-likelihood term in equation 1 and then compute the KL divergence between the agent’s weight priors and its sampled weight values (for both the actor and the critic), resulting in the following loss function.

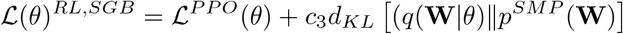

Where *c*_3_ is a further hyperparameter controlling the relative weighting of the PPO loss to the KL divergence.

### 4.4 Supervised Learning Hyperparameters

#### 4.4.1 MNIST

For MNIST experiments, we used 3-layer feedforward networks with 800 hidden units. To obtain SGB’s performance on MNIST across different compression ratios, we fixed the number of cell types in the input and output layer to be 30 and 10 respectively, and varied the number of cell types in the hidden layer. A list of hyperparameters used to train the SGB for the feedforward networks on the MNIST task has been summarized in Table 1.

**Table 1:** Hyperparameters for SGB for CIFAR and MNIST Experiments.

| Hyperparameter | MNIST | CIFAR |
| --- | --- | --- |
| Learning Rate | 0.00005 | 0.001 |
| Batch Size | 64 | 128 |
| Number of Epochs | 50 | 200 |
| Optimizer | Adam | Adam |

**Table 2:** Hyperparameters for Brax experiments.

| Hyperparameter | Ant | Halfcheetah |
| --- | --- | --- |
| Learning Rate | 0.0003 | 0.0003 |
| PPO clip coefficient | 0.001 | 0.3 |
| Number of parallel envs | 2048 | 2048 |
| Entropy coefficient | 0.01 | 0.001 |

**Table 3:**
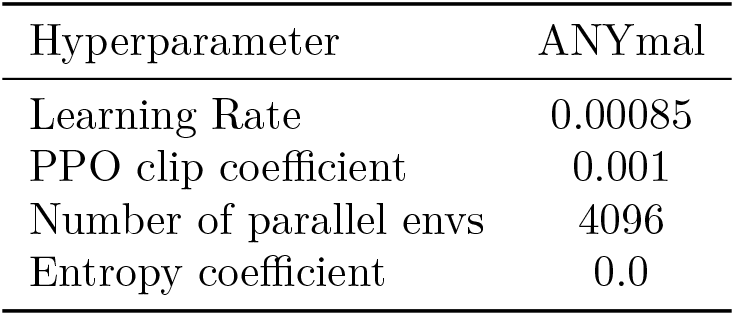
Hyperparameters for ANYmal experiments.

| Hyperparameter | ANYmal |
| --- | --- |
| Learning Rate | 0.00085 |
| PPO clip coefficient | 0.001 |
| Number of parallel envs | 4096 |
| Entropy coefficient | 0.0 |

#### 4.4.2 CIFAR

For the CIFAR experiments, we used a 5-layer CNN based architecture consisting of two initial convolutional layers followed by three fully connected layers. To ensure consistency in experimentation we only compressed the fully connected layers in the network. To obtain SGB’s performance on CIFAR across different compression ratios, we fixed the number of cell types in the first fully connected layer and output layer to be 100 and 10 respectively, and varied the number of cell types in the hidden layer. A list of hyperparameters used to train the SGB on the CIFAR task has been summarized in Table 1.

### 4.5 RL Experiment Hyperparameters

For our Brax experiments we use the Brax implementation of PPO (Freeman et al., 2021), while for the robotic control experiments we use the cleanRL PPO implementation (Huang et al., 2022). Below we briefly list hyperparameters we found to be important for performance, the full set of hyperparameters are in the supplementary code.

## A Proofs

### A.1 Notation

Let us denote the loss function as ℒ and the loss value at parameters *w* as ℒ (*w*). Let us denote the parameters of the Bayesian network as (*µ, σ*) where *µ* and *σ* are vectors indicating the mean and variance respectively of the distribution of each parameter of the network. In a vanilla setting (no compression), each element of (*µ, σ*) corresponds to the distribution for a different parameter in the network. Let the network have *P* parameters. So, *µ, σ*∈ ℝ^*P*^. We will also use ∇ℒ (*w*) and ∇^2^ℒ (*w*) to denote the gradient and Hessian respectively of the loss function ℒ at parameter values *w*. Given these notations, the parameters of a sampled network can be written as *w* = (*µ* + *σ*⊙ *ϵ*), where *ϵ* is a vector with elements drawn from a standard Gaussian distribution.

### A.2 Key Theoretical Results

In this section, we provide an analytical framework to elucidate the advantages of stochasticity during development, with particular focus on explaining the empirical results in subsection 2.4. First, we show that the difference between the mean performance of sampled networks and the performance of the mean network depends on the variance in the SGB network parameters and the eigenvalues of the loss function Hessian. Next, we study the implicit bias of gradient descent in modulating this difference between the performances. Finally, we posit that the SGB network parameters would converge to saddle points in the loss landscape during optimization, thereby leading to a better mean performance of sampled networks, compared to the performance of the mean network.

Let us denote the network parameter space as ℝ^*p*^, i.e. a *p*-dimensional space, the task-relevant loss function asℒ : ℝ^*p*^ →ℝ, the gradient of the loss function as ∇ℒ: ℝ^*p*^→ ℝ^*p*^ and the Hessian as ∇^2^ℒ : ℝ^*p*^→ ℝ^*p*×*p*^. The parameters of a sampled network are denoted as *θ* ∈ ℝ^*p*^, while the parameters for the mean and variance of the Bayesian network are denoted as *µ*∈ ℝ^*p*^ and *σ* ∈ ℝ^*p*^, respectively. Under this notation, we can write the parameters of a sampled network, *θ* = *µ* + *σ ϵ*, where *ϵ* ∈ ℝ^*p*^ is a vector with each element drawn i.i.d. from a unit Gaussian. Note that this is the simplest form of Bayesian neural network parameterization, i.e. without any compression. This framework can be easily extended to networks with a compression ratio more than 1 by enabling parameter sharing such that *µ*, 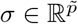, where 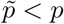.

#### Theorem A.1.

*The difference between the mean loss of sampled networks*, 𝔼_*ϵ*_ [ℒ (*µ* + *σ* ⊙ *ϵ*)] *and the loss of the mean network (i*.*e. network with parameters, θ* = *µ)*, ℒ (*µ*) *is as follows*.

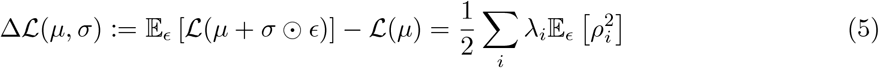

*where* (*λ*_*i*_, *ν*_*i*_)*’s denote the eigenvalue and eigenvector pairs of* ∇^2^ℒ *and ρ*_*i*_ = (*σ* ⊙ *ϵ*)^*T*^ *ν*_*i*_.

A direct corollary of theorem A.1 is that for convex loss landscapes, i.e. when *λ*_*i*_ *>* 0 ∀*i*∈ {1 … *p* }, 𝔼_*ϵ*_ [ℒ (*µ* + *σ* ⊙ *ϵ*)] *>ℒ* (*µ*). This would mean that the mean network would actually perform better than the average sampled network. But, given our empirical observations, we posit that the optimization loss landscape is non-convex for the sorts of tasks that animals must solve. Next, we consider the implicit bias of gradient descent during optimization of the Bayesian neural network parameters, *µ* and *σ*, within the SGB framework.

#### Theorem A.2.

*Let* 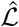 *denote the variational inference loss (computed using the reparameterization trick) for training the Bayesian neural network parameters µ, σ. The parameter updates following a gradient descent step on* 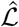 *are as follows*.

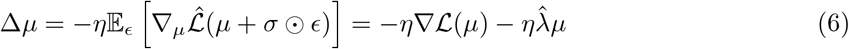

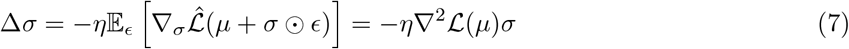

*where η is the learning rate and* 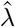 *is the regularization weight for the KL divergence term in the variational inference (ELBO) loss, representing the weight for the prior*.

Note that theorem A.2 implies that change in *µ* is akin to doing gradient descent on a single network with parameters *µ*. Thus, we can extrapolate the behavior of gradient descent (and its stochastic versions) in high-dimensional non-convex loss landscapes to the SGB setting and hypothesize that the mean parameters will converge to saddle points in the loss landscape (Chen et al., 2024). Another direct consequence of theorem A.2 is that the modulation of *σ* is controlled by the eigenvalues of the Hessian of the ℒ. Specifically, the projection of *σ* along the directions corresponding to positive (negative) eigenvalues decreases (increases). Consequently, the 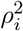 values corresponding to positive (negative) eigenvalues decrease (increase). Combining this result with theorem A.1, we can infer that Δ ℒ (*µ, σ*) decreases after a step of gradient descent. Over the course of optimization, the parameters of SGB will converge to regions in the (non-convex) loss landscape such that Δ ℒ (*µ, σ*) *<* 0, i.e. the mean loss of the sampled networks is less than the loss of the mean network. Assuming a monotonic relationship between the in-distribution performance performance and training loss (whereby higher performance corresponds to lower loss), we have the desired result. Thus, our empirical observation that the average sampled network outperforms the mean network is analytically grounded, as long as the loss is non-convex.

### A.3 Effect of sampling on the Loss

#### Theorem A.3.

*The difference between the mean loss of sampled networks, denoted as* 𝔼_*ϵ*_ [ℒ (*µ* + *σ* ⊙ *ϵ*)] *and the loss of the mean network (i*.*e. network with parameters, θ* = *µ), denoted as* ℒ (*µ*) *is as follows*.

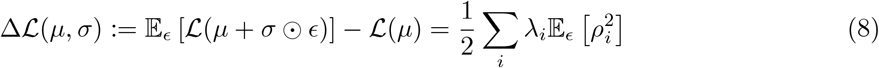

*where* (*λ*_*i*_, *ν*_*i*_)*’s denote the eigenvalue and eigenvector pairs of* ∇^2^ℒ *and ρ*_*i*_ = (*σ* ⊙ *ϵ*)^*T*^ *ν*_*i*_.

*Proof*. We will start by using Taylor series expansion of the loss function. Let us write the loss function for a sampled network with parameters *w* = (*µ* + *σ*⊙ *ϵ*) using a Taylor Series expansion around *w* = *µ*:

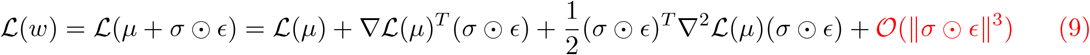

Assuming that 3rd (and higher) order terms are significantly smaller, we can drop the terms in red. Next, taking an expectation over different *ϵ*:

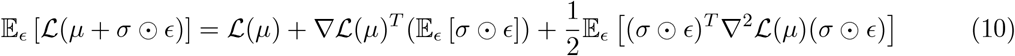

Note that 𝔼_*ϵ*_ [*ϵ*] = 0 and consequently 𝔼_*ϵ*_ [*σ*⊙ *ϵ*] = 0. Therefore, the first order term disappears. To simplify the second order term, we can assume (without loss of generality) that ∇ℒ^2^ (*µ*) has eigenpairs { (*λ*_1_, *ν*_1_), (*λ*_2_, *ν*_2_) … (*λ*_*P*_, *ν*_*P*_) } such that {*ν*_1_, *ν*_2_ … *ν*_*P*_} are a set of orthonormal basis vectors in the P-dimensional parameter space. Note that all (*λ, ν*) pairs are functions of *µ*, but we have dropped the explicit notation for sake of clarity. We can use this orthonormal basis set to write (*σ* ⊙*ϵ*) as a linear combination of *ν*_*i*_’s. Specifically,

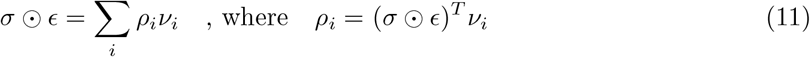

It is worth noting that: 𝔼_*ϵ*_ [*ρ*_*i*_] = 𝔼_*ϵ*_ [(*σ*⊙ *ϵ*)]^*T*^ *ν*_*i*_ = 0 We can now use this linear combination to simplify the quadratic term:

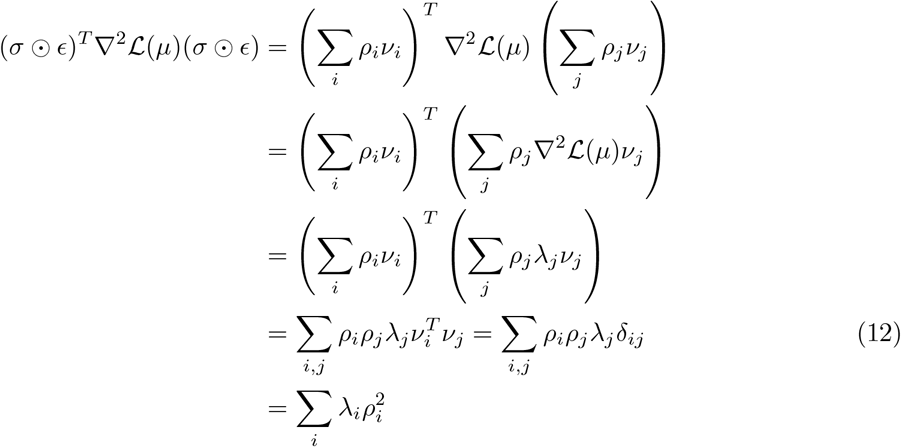

Above, in eq. (12), we have used the fact that 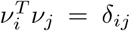, where *δ*_*ij*_ indicates the Dirac-delta function, i.e. it is 1 when *i* = *j* and 0 otherwise. This fact is a direct consequence of (*ν*_*i*_, *ν*_*j*_) being part of the orthonormal basis set.

Plugging these results about the first and second order terms back in eq. (10):

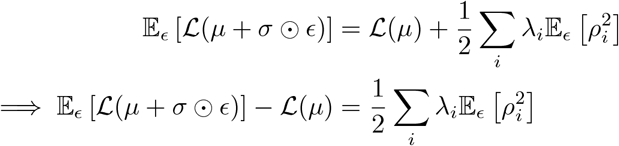

**Key takeaways:** The above equation describes the difference between the mean loss of the population and the loss of the mean of the population. Note that 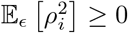.

1. In a strictly convex setting, i.e. *λ*_*i*_≥ 0 ∀*i*, the right hand side is strictly non-negative. Thus, the mean loss (performance) of the population will be more (less) than the loss (performance) of the population mean network.
2. In a non-convex setting, i.e. *λ*_*i*_ *<* 0 for some *i*, it is possible to have the right hand side of the equation to be negative. Thus, it is possible to have a certain variance structure in the network parameter space (*σ*) such that the mean loss (performance) of the population will be less (more) than the loss (performance) of the population mean network.

### A.4 Effect of gradient descent on Bayesian network parameters

#### A.4.1 Background: variational inference with Bayesian neural networks

**Notation:** Let us introduce a few more notations to denote the dataset as 𝒟 which consists of a set of inputs and corresponding labels: { (*x*_1_, *y*_1_), (*x*_2_, *y*_2_) … (*x*_*i*_, *y*_*i*_) … (*x*_*n*_, *y*_*n*_) }. We present a supervised learning problem here, but this notation can be extended to unsupervised, self-supervised as well as reinforcement learning setups. The task is to learn network parameters *w* such that the network’s predictions, denoted by *ŷ*(*x*_*i*_) = *f* (*x*_*i*_, *w*) are close to *y*_*i*_. We can formulate this objective as a reducing the negative log likelihood of the observed *y*_*i*_ to be drawn from the distribution *p*(*ŷ*| *x*_*i*_, *w*) which depends on the network’s parameters.

**Variational Inference perspective:** We can use a variational inference perspective to solve the above task, wherein the problem is to learn a distribution of network parameters, denoted by *q*_(*µ,σ*)_(*w*), which best matches the posterior distribution of the network parameters given the observed data, *p*(*w*|𝒟). For the Bayesian network perspective, we can assume that *q*_(*µ,σ*)_(*w*) has the following Gaussian form:

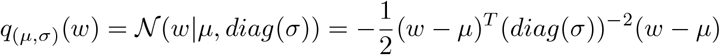

where *diag*(*σ*) denotes a diagonal matrix with elements of *σ* in the diagonal.

The true objective is to reduce the KL-divergence between *q*_(*µ,σ*)_(*w*) and *p*(*w*|𝒟).

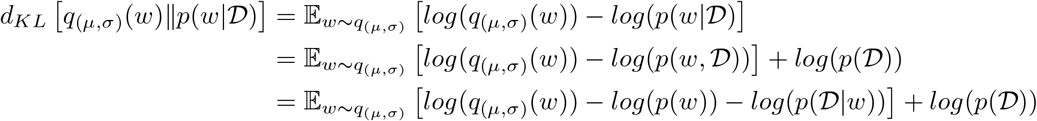

Here, *p*(*w*) is the prior over network parameters. We can also assume this to be have a Gaussian form with mean 0:

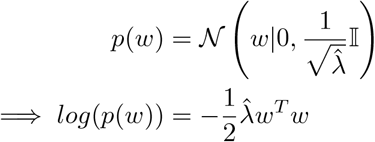

where 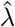 can be thought of as the weight decay hyperparameter, or how strongly we want to rely on the chosen prior of a mean 0 distribution.

Based on these expressions, we can define the following loss:

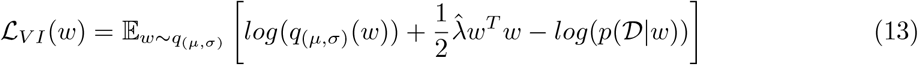

Minimizing this loss will lead to minimizing the overall objective of reducing the KL-divergence. However, evaluating and minimizing this loss requires us to sample from the current estimate of the posterior distribution and then backpropagate gradients through the distribution parameters, which if often intractable.

##### Reparameterization trick

To make the problem tractable, we will use the reparameterization trick, wherein we will write *w* = *µ*+*σ*⊙ *ϵ*, wherein elements of *ϵ* are drawn from a standard Gaussian. In doing so, the *log*(*q*_(*µ,σ*)_(*w*)) becomes a constant. Note that this is exactly the Bayesian neural network framework, as described above! Now, we can write a tractable version of eq. (13):

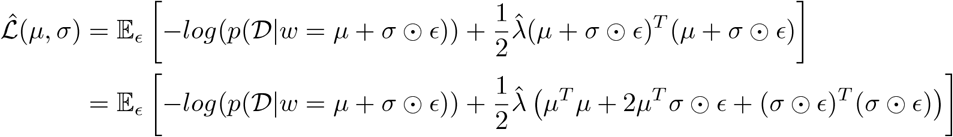

Note that the first term is the negative log likelihood of the data, given the current network parameters. Therefore, we can represent it as the task loss, ℒ (*w* = *µ* + *σ* ⊙ *ϵ*). Thus,

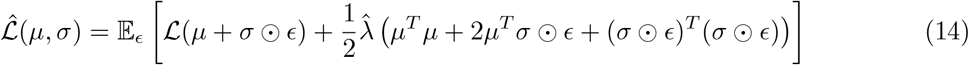

#### A.4.2 Impact of gradient descent

We can now introduce the theorem from the main text and present its proof.

##### Theorem A.4.

*Let* 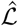 *denote the variational inference loss, computed with reparameterization trick, that is used to train the Bayesian neural network parameters µ, σ. The updates to the parameters following a gradient descent step on* 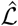 *are as follows*.

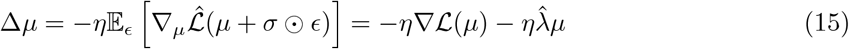

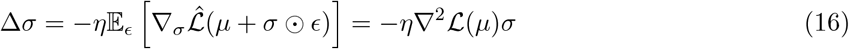

*where η is the learning rate and* 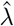 *is the weight for the KL divergence term in the variational inference (ELBO) loss, i*.*e. the weight for the prior*.

*Proof*. Let us write the gradients with respect to the mean and variance parameters of our distribution. Before we dive in to deriving the expressions for the parameter updates, we present some results that are consequences on the Chain rule and will be useful.

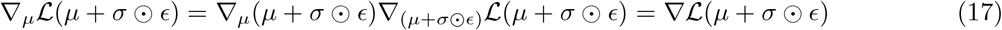

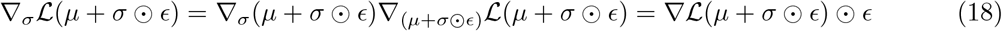

Focusing on the expression for mean, *µ*:

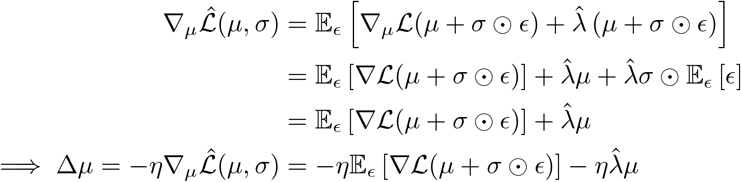

where *η* is the learning rate. Again, we can use the Taylor Series expansion for∇ℒ (*µ* + *σ*⊙ *ϵ*) around *w* = *µ*:

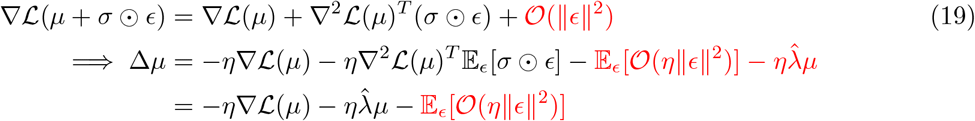

Ignoring the third (and higher) order terms of *η* and ∥*ϵ*∥,

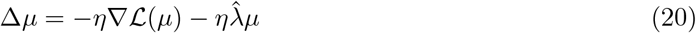

Now, we can repeat the same process for variance, *σ*.

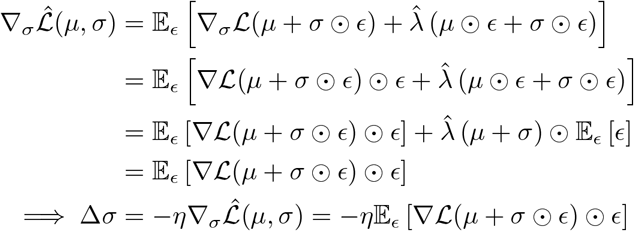

Using the Taylor Series expansion for ∇L(*µ* + *σ* ⊙ *ϵ*) around *w* = *µ* from eq. (19):

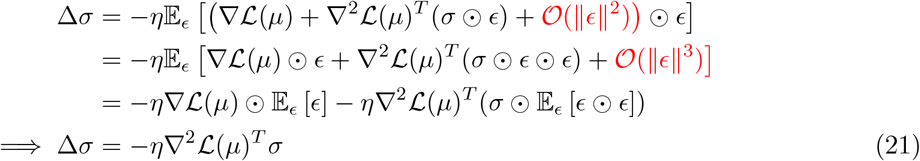

Without loss of generality, we can assume that the Hessian is symmetric (the order of derivatives can be interchanged). So,

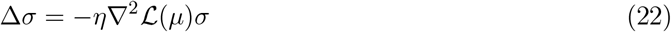

### A.5 Intuitive explanation of the consequences of above Theorems

Let us write *σ* using the basis set of (*ν*_1_, *ν*_2_ … *ν*_*P*_), i.e. the eigenvectors of ∇ℒ^2^ (*µ*), to further simplify the expression of Δ*σ* in theorem A.4.

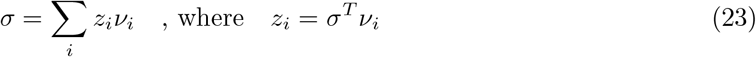

Note that *ν*_*i*_’s are dependent on *µ* and can be considered to be constant while updating *σ*. So,

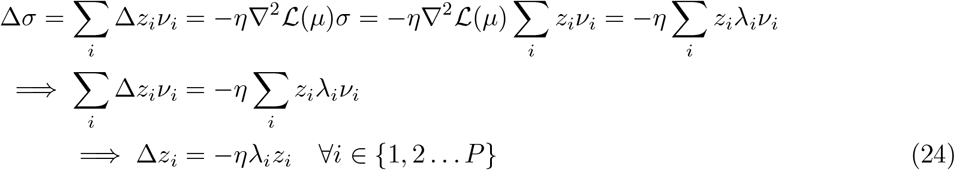

This equation indicates that directions corresponding to *λ*_*i*_ *>* 0 would correspond to 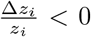 and consequently lead to pushing *z*_*i*_ closer to 0, i.e. shrink variance in those directions. The contrary will be true in directions corresponding to *λ*_*i*_ *<* 0, i.e. variance will expand in those directions. Thus, gradient descent leads to a *reduction* of the term in theorem A.3 that quantifies the difference between the mean loss of the population and the loss of the population mean.

#### Key takeaways

1. The changes in the mean are like doing gradient descent on a neural network with the parameters equal to the mean.
2. The changes in the variance are such that the variance along strictly convex directions are reduced and variance along strictly concave directions are increased.

Taken together, we have presented theoretical justification that leverages the effect of gradient descent and sampling in Bayesian neural networks to explain why the mean performance of the population can be better than the performance of the population mean network.

## B Real Environment Interfaces for RL tasks

In this section, we provide a detailed view of the real environment interfaces used for the reinforcement learning (RL) tasks discussed in the main text. Specifically, we focused on the motor control tasks involving the Ant, HalfCheetah, and Anymal environments, which are part of the Brax framework. The Ant and HalfCheetah tasks require the agent to learn efficient locomotion strategies, balancing speed and stability, while navigating diverse terrains. The Anymal environment, a simulated quadruped robot, introduces additional complexity with varied terrains, making it an excellent testbed for assessing the ability of RL models to generalize across different physical conditions.

## Footnotes

1 Adapted from (Azabou et al., 2023)

